# Application of large language models to the annotation of cell lines and mouse strains in genomics data

**DOI:** 10.64898/2026.03.05.709906

**Authors:** Sanja Rogic, B. Ogan Mancarci, Brianna Xu, Anna Xiao, Carlton Yan, Paul Pavlidis

## Abstract

Accurate, consistent and comprehensive metadata are essential for the reuse of functional genomics data deposited in repositories such as the Gene Expression Omnibus (GEO), however, achieving this often requires careful manual curation that is time-consuming, costly and prone to errors. In this paper, we evaluate the performance of Large Language Models (LLMs), specifically OpenAI’s GPT-4o, as an assistive tool for entity-to-ontology annotation of two commonly encountered descriptors in transcriptomic experiments - mouse strains and cell lines. Using over 9,000 manually curated experiments from the Gemma database and over 5,000 associated journal articles, we assess the model’s ability to identify relevant free-text entries and map them to appropriate ontology terms. Using zero-shot prompting and retrieval-augmented generation (RAG) to incorporate domain-specific ontology knowledge, GPT-4o correctly annotated 77% of mouse strain and 59% of cell line experiments, and uncovered manual curation errors in Gemma for over 200 experiments. GPT-4o substantially outperformed a regular expression–based string-matching method, which correctly annotated only 6% of mouse strain experiments due to low precision. Model errors often arose from typographical mistakes or inconsistent naming in the GEO record or publication, and resembled those made by human curators. Along with annotations, our approach requests that the model output supporting context and quotes from the sources. These were typically accurate and enabled rapid curator verification. These findings suggest that LLMs are not ready to fully replace manual curators, but can already effectively support them. A human-in-the-loop workflow, in which LLM’s annotations are provided to human curators for validation, may improve the efficiency and quality of large-scale biomedical metadata curation.

## Introduction

Biomedical databases and repositories are essential for gathering and organizing the vast amounts of generated data, ensuring that these resources are findable and accessible to the scientific community. One such repository is the NCBI Gene Expression Omnibus (GEO), which has accumulated hundreds of thousands of high-throughput functional genomic datasets spanning diverse technologies, organisms, tissues and experimental conditions (Clough et al., 2023). These data are consistently used for secondary analysis, meta-analysis, method development and development of derivative databases. However, the re-use of the data critically depends on the availability of accurate and consistent metadata. While GEO has established submission formats and metadata standards, these are intentionally flexible and not strictly enforced to reduce the burden on data submitters. This can lead to missing, ambiguous or mistyped experiment information, discrepancies with the associated publication and broader inconsistency across studies.

To make GEO datasets more findable, comparable and re-usable, resources like Gemma (Lim et al., 2021) curate experiments and map free-text descriptions onto controlled vocabularies and ontologies, including terms for features such as organisms, strains, cell lines, drugs, genotypes, tissues, and diseases. Ontology-based annotation enables more effective database querying through ontology inference, facilitates data aggregation across studies, and simplifies downstream computational analyses. However, manual curation is resource-intensive and does not eliminate the risk of errors.

Large language models (LLMs) offer a potential way to expedite and facilitate the curation process due to their ability to interpret natural-language inputs and find key concepts within complex writing. Unlike approaches that rely primarily on string similarity and predefined lexicons, LLMs use contextual cues to disambiguate entities. Recent models such as GPT-4o can process long heterogeneous inputs and follow complex instructions, making them good candidates for data curation tasks. Previous work showed promising results for a range of data annotation and information extraction tasks (Chen et al., 2025; Gue et al., 2024; Jensen et al., 2025; Kainer, 2025; Mahmoudi et al., 2024; Poretsky et al., 2025; Riquelme-García et al., 2025; Turner et al., 2025), but these studies were typically limited in scope, either relying on sentence-level inputs or evaluating performance on only a few dozen scientific articles.

In this study, we evaluate the ability of GPT-4o to annotate two types of entities that are often encountered in metadata of transcriptomic experiments: mouse strains and cell lines. Using over 9,000 manually curated experiments from the Gemma database as a reference, as well as over 5,000 associated journal articles, we assess GPT-4o’s performance in identifying the relevant free-text entries from GEO metadata and linked publications and mapping them to domain-specific ontology terms. We employ a zero-shot prompting strategy and use a retrieval-augmented generation (RAG) approach to ground the model with pre-compiled lists of ontology terms. For comparison, for the mouse strain task we also evaluate a regular expression–based string-matching approach. These findings provide an initial, systematic assessment of GPT-4o as an assistive tool for ontology-based curation of transcriptomic experiments and highlight both its promise and current limitations of LLMs for large-scale biomedical metadata annotation.

## Methods

### Compiling domain-specific knowledge

To provide GPT-4o with relevant external domain-specific knowledge, we compiled separate lists of ontology terms for each task. For the mouse strain identification task, we used the Experimental Factor Ontology (EFO) and the Gemma Ontology (TGEMO) to extract all child terms (i.e., mouse strains) of the term “Mus musculus” (http://purl.obolibrary.org/obo/NCBITaxon_10090). This resulted in a list of 156 commonly used mouse strains.

For the cell lines identification task, we combined all terms from the Cell Line Ontology (CLO) with the child terms of “cell” from EFO (http://purl.obolibrary.org/obo/CL_0000000). After removing duplicates, the final list included 46,032 unique cell lines.

### Prompt design

For each transcriptomic experiment in the test dataset, we generated a GPT-4o prompt, beginning with a plain-text description of the task, followed by the experiment’s metadata sourced from GEO in JSON format. The metadata included the experiment title, summary and overall design, as well as characteristics and protocol descriptions for each individual sample. If the experiment has an associated publication that is publicly accessible through PubMed Central, we appended its title, abstract and Methods section. For the mouse strain identification task, the prompt also included labels, URIs and descriptions of compiled ontology terms, formatted as a JSON object. Finally, GPT-4o was instructed to format its output according to a provided JSON schema to enable programmatic processing.

### Running experiments

For all our analyses, we used OpenAI’s GPT-4o model (gpt-4o-2024-11-20) via its API and the OpenAI Python library (version 1.54.1). To make our results more reproducible, the temperature parameter was set to 0 and the seed was set to 1.

For the mouse strain identification task, GPT-4o was prompted to return both the labels and URIs of all mouse strains detected in the input, along with verbatim quotes supporting each decision (Figure 1a).

**Figure 1.**
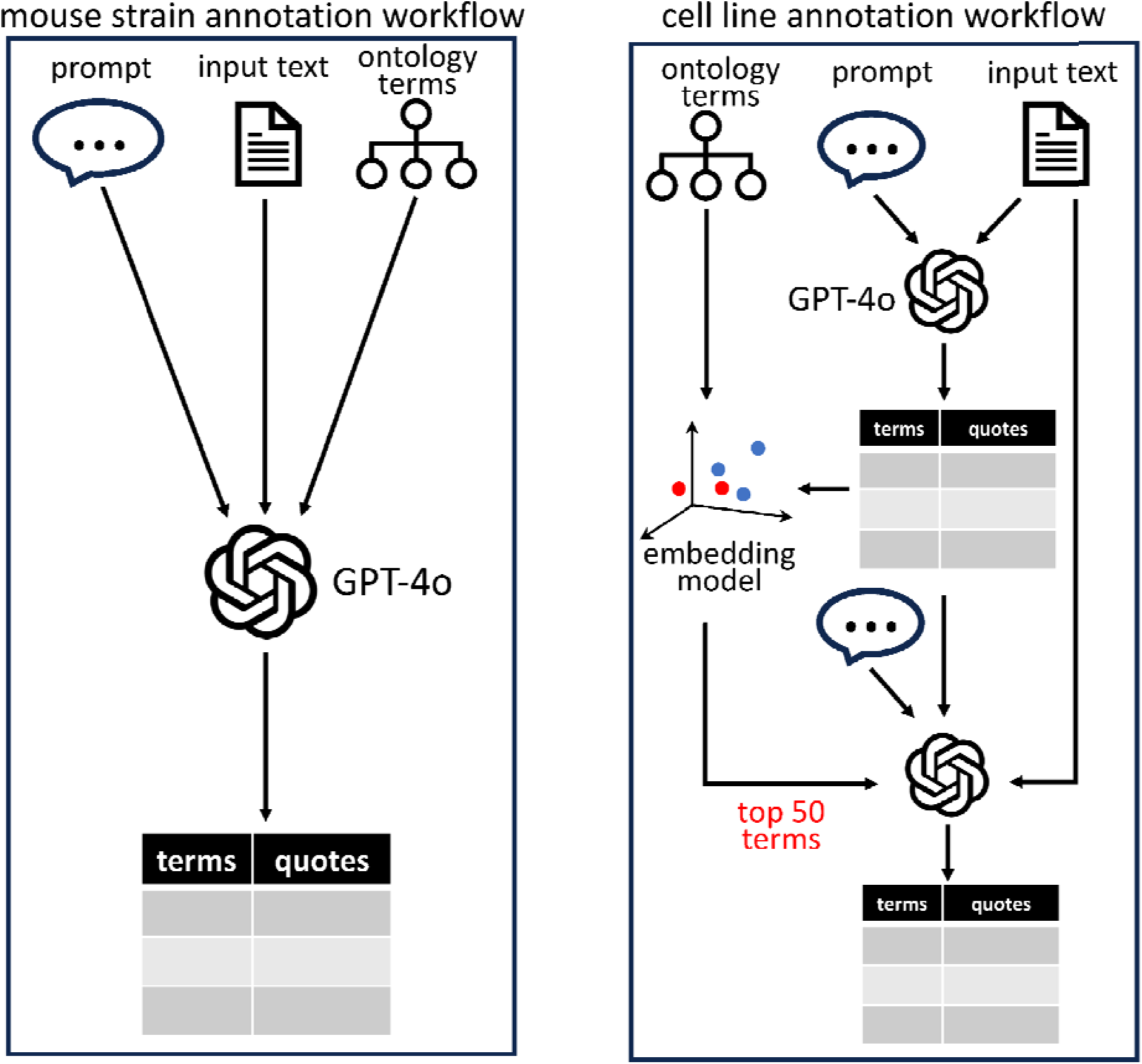
Curation workflows for the two types of annotation tasks. Each workflow begins with a prompt and input text consisting of the experiment’s metadata from GEO (title, summary, overall design, sample-level characteristics and protocol descriptions) and, when available, associated publication (title, abstract and Methods sections). a) The input for the mouse strain task also includes a list of ontology terms (labels, URIs and descriptions). GPT-4o is prompted to return the labels and URIs of all mouse strains detected in the input, along with verbatim quotes supporting each decision. b) Annotation of cell lines is a two-step process. First, GPT-4o is prompted to return cell lines identified in the input as free-text entries, together with supporting quotes. Next, these responses are embedded and compared against a pre-generated vector database of cell line ontology terms, and the top 50 most similar terms are returned to GPT-4o along with the original input text and first-stage outputs. Finally, GPT-4o is instructed to select the corresponding ontology term.

Due to GPT-4o’s limited context length (128k tokens), we were unable to include the full list of 46,032 cell line ontology terms in a single prompt. Instead, the terms and their descriptors were embedded using the “text-embedding-3-large” model to create a vector database of cell line embeddings. Annotation using GPT-4o then proceeded in two stages (Figure 1b). First, GPT-4o was asked to identify cell lines mentioned in the input text and to return them as free-text entries, along with verbatim quotes supporting each identification. Next, these free-text responses were embedded and compared against the previously generated ontology term vector database. For each response, the top 50 most similar ontology terms were then returned to GPT-4o, along with the original input text and first-stage outputs, which was then instructed to replace the free-text entries with the corresponding specific ontology terms.

### Annotation using string matching

To compare GPT-4o’s performance against a simpler computational approach, we used a string-matching method based on regular expressions to search for ontology terms or their synonyms from the task-specific ontology lists within the same input text provided to GPT-4o. To minimize random or spurious matches, we excluded all terms shorter than three characters. We also removed terms that commonly appear as ordinary English words in the text, such as NOR (used for the NOR mouse strain) or CAST (a synonym for the CAST/EiJ mouse strain). An annotation was assigned to an experiment when a case-insensitive match was detected.

The output was evaluated using the same criteria as for the GPT-4o’s evaluation. However, mismatches with Gemma’s annotation generated by this method were not manually reviewed by the curators, due to the large number of erroneous annotations produced.

### Experiment curation in Gemma

The Gemma manual curation procedure is described in Lim et al. (2021). We provide a brief overview here for context. The process in Gemma begins by automatically parsing and uploading the experiment’s metadata from GEO SOFT files. All experiments are then subjected to careful manual curation performed by trained undergraduate research assistants who follow an established set of guidelines designed to ensure consistency across datasets. The curation process captures both the experimental design and the scientific topics of each study by mapping free-text metadata and publication descriptions to controlled vocabulary terms.

Gemma uses ontology-based annotation to assign category–value pairs to relevant biological entities in the metadata. Examples of categories include tissue or organ, disease, treatment, sex, mouse strain and cell line. Gemma supports several established biomedical ontologies for annotating values within these categories, of which EFO for mouse strains and CLO for cell lines are most relevant to this paper. Built-in curation tools assist curators in selecting appropriate ontology terms by suggesting previously used terms and encouraging consistent usage. Annotations may be reviewed by a second curator and periodically audited to ensure quality and consistency.

### Evaluation of automatic annotations

We first performed an automated comparison between the annotations produced by GPT-4o or the string-matching baseline and the existing Gemma annotations. Experiments for which the predicted ontology terms exactly matched the corresponding Gemma terms were classified as correctly annotated. For all remaining experiments, curators manually reviewed discrepancies to determine the source of the mismatch and, when necessary, corrected Gemma annotations before finalizing their assessment.

To quantify predicted annotation accuracy, for each experiment, we computed the following metrics:

- True Positives (TP) – number of correctly predicted ontology terms
- False Positives (FP) - number of incorrectly predicted ontology terms, i.e., not present in Gemma’s annotation
- False Negatives (FN) - number of ontology terms present in Gemma’s annotation that were missed by the evaluated approach

Based on these metrics, we computed the following performance measures for each experiment:

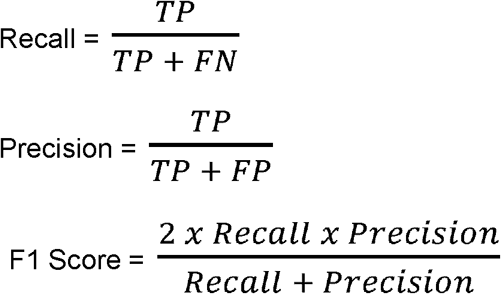

To ensure meaningful estimates, experiments without any annotations in Gemma (TP + FN = 0) were omitted from the computation of average recall, while experiments for which the evaluated approach produced no annotations (TP + FP = 0) were omitted from the computation of average precision.

## Results

We conducted all experiments using GPT-4o (released in May 2024), the most advanced OpenAI model available during the study period. Using the model’s API we annotated two types of information commonly found in GEO’s metadata for transcriptomic experiments: mouse strains and cell lines. Performance was evaluated using experiments from Gemma, which have been manually annotated with ontology terms from these categories. Gemma’s data curation and annotation are performed by trained undergraduate research assistants, and various internal checks and reviews are used to ensure high accuracy; however, errors can occur.

For each experiment, we constructed a prompt that included the GEO metadata and, when available, the Methods section of the associated publication. No examples of correct annotations were provided, consistent with the zero-shot prompting approach. We applied a Retrieval-Augmented Generation (RAG) approach to ground the LLM with domain-specific knowledge, using pre-compiled lists of ontology terms relevant to each task. To facilitate evaluation, GPT-4o was instructed to produce programmatically consumable output along with supporting evidence for each annotation (Figure 2).

**Figure 2.**
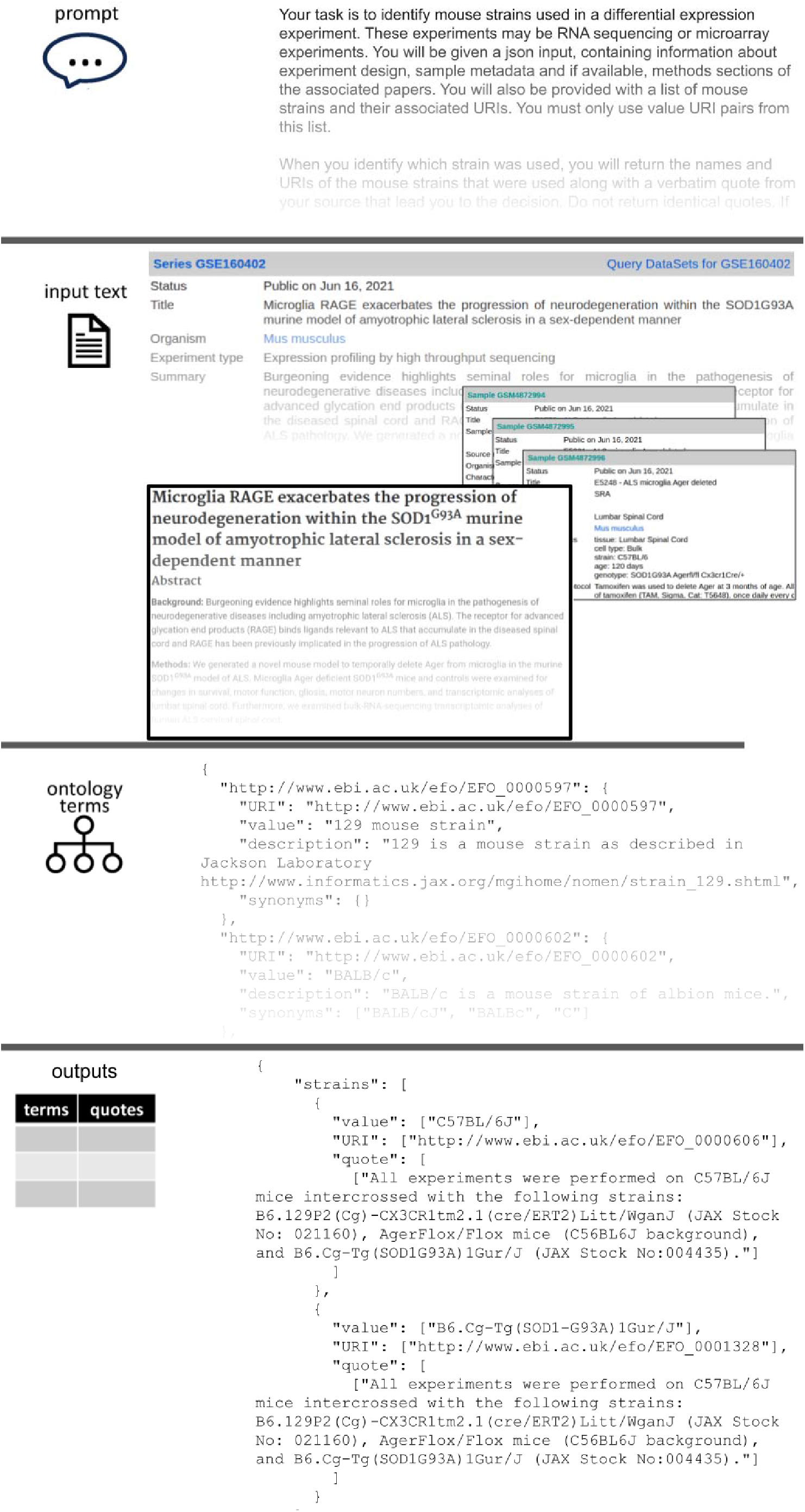
Example of the mouse strain annotation workflow for GEO experiment GSE160402, including the prompt, input text, ontology term list (in JSON format), and the resulting output (in JSON format).

### Annotation of mouse strains

For the mouse strain annotation task, we selected a set of previously curated transcriptomic experiments from Gemma, using the existing Gemma annotations as the baseline. We included experiments annotated with at least one ontology term from the task-specific lists described above, yielding 6,013 experiments in total, 3,362 of which had associated publications (Table 1). The automated comparison between GPT-4o and Gemma annotations yielded 4,214 exact matches. Manual inspection of the remaining 1,799 experiments identified an additional 422 cases in which GPT-4o correctly captured all relevant mouse strains, bringing the total number of correctly annotated studies to 4,636 (77%). Manual review also revealed 230 studies in which Gemma’s annotations were inaccurate; these were corrected prior to completing the evaluation. Among the 1,377 studies where GPT-4o’s output still disagreed with the corrected Gemma annotations, most discrepancies were due to GPT-4o missing some strains (1,286), while in 893 cases it also added incorrect annotations. Overall, average (± standard deviation) recall was 0.82±0.36 and average precision 0.82±0.36. The mean F1 score was 0.82±0.36.

**Table 1:** Summary of task-specific evaluation datasets and corresponding performance results.

|  | Experiments analyzed (with publication) | Manually reviewed experiments | Accurately annotated experiments | Experiments without false negatives | Experiments without false positives | Mean recall | Mean precision | Corrections of existing annotations |
| --- | --- | --- | --- | --- | --- | --- | --- | --- |
| Mouse strain annotations | 6,013 (3,362) | 1,799 | 4,636 (77%) | 4,727 (79%) | 5,120 (85%) | 0.82 | 0.82 | 230 |
| Cell line annotations | 3,377 (1,868) | 1,063 | 1,990 (59%) | 2,083 (62%) | 2,215 (65%) | 0.72 | 0.72 | 17 |

### Annotation of mouse strains using string matching

To assess GPT-4o’s performance relative to a simpler method, we used regular expression– based string matching to identify mouse strains (this was not computationally feasible with the cell line task due to the large number of terms). While this approach achieves relatively high recall, detecting all relevant terms in 74% of cases, it also produces false positive annotations in 80% of experiments. Only 6% of experiments were annotated correctly overall. This corresponds to an average recall of 0.70±0.46, an average precision of 0.32±0.27 and an average F1 score of 0.42±0.30.

### Annotation of cell lines

GPT-4o’s performance on the cell line annotation task was evaluated using a similar procedure. Among 3,377 evaluated studies (1,868 with associated publications), GPT-4o produced perfect annotations for 1,990 of them (59%) (Table 1). Of the remaining 1,387 studies, 1,294 contained spurious annotations and 1,162 had missed annotations. The per-study accuracy metrics were: mean recall = 0.72 ± 0.43, mean precision = 0.72 ± 0.43, and mean F1 score = 0.72 ± 0.43. The lower accuracy compared to the strain annotation task likely reflects the substantially larger cell line ontology dictionary (46,032 terms) and high lexical complexity of the cell line ontology labels.

### Detection of human curation errors

GPT-4o’s output helped correct existing Gemma annotations in 230 studies for mouse strains and 17 for cell lines. For mouse strains, most curation errors stemmed from selecting the incorrect version among closely related strains, often due to inconsistent reporting within GEO metadata and/or the associated publication. For example, the GEO sample-level metadata for experiment GSE135602 specifies the mouse strain as FVB. However, the Methods section of the corresponding paper (Herrero-Navarro et al., 2021) identifies the strain more precisely as FVB/N, which our curators missed. Similar inconsistencies can occur within different sections of GEO metadata: for instance, the record of GSE127190 lists C57BL/6J in its overall design, but C57BL/6 in its sample metadata. Overall, at least 37% of the cases had such inconsistencies. Although improvements in curator training and curation procedures could potentially mitigate such issues, GPT-4o’s ability to quickly analyze the entirety of provided information proved especially effective in identifying and correcting these inconsistencies.

### Characterizing GPT-4o annotation errors

A closer examination of GPT-4o’s errors revealed that they were often similar in nature to those made by human curators. Many arose from typographical errors or alternative spellings in GEO metadata or in the associated publication. For instance, GPT-4o failed to return the mouse strain C57BL/6 for dataset GSE55188 because it was incorrectly referenced by the submitter as “C57/Bl6”. Similarly, for dataset GSE56690 the model returned C57BL instead of C57BL/6, most likely due to the misplaced slash in the strain name in the GEO record (C57/BL6). In general, we observed that GPT-4o was more likely to make mistakes in experiments where string matching failed to detect the correct strains than when it succeeded (25.6% vs 20.2% error rate; one-sided Fisher’s exact p = 6.5 × 10□□), suggesting that these cases present challenges even for context-aware models.

We also observed some hallucinatory errors, where the model produces terms not present in the input text; for example, in dataset GSE59018, GPT-4o reported FVB/NJ despite the only strain mentioned in the input text being FVB. Notably, however, in such cases the supporting quotes returned by the model were consistently faithful to the source text and always contained the correct term. This allows a human curator to quickly identify the error and make the necessary correction.

Using the two-step querying approach for cell lines allowed us to work with a much larger list of ontology terms, but it also introduced additional sources of error in the annotation process. We observed failures in both steps: either the model failed to identify valid cell line mentions in the input text or the correct ontology term was not retrieved among the top 50 candidate terms from the embedding-based similarity search. Errors in the first stage typically arose because entity identification was performed without ontology-constrained candidate terms. For example, for dataset GSE1977 GPT-4o mistakenly identified “NoCa” as a cell line name, which the authors used as a designation for patients with no cancer. Dataset GSE41790 is an example of failure in the second stage, where the correct cell line, “BJ cells”, was identified initially, but the corresponding ontology term was ranked 87th based on the similarity search and therefore not provided to GPT-4o for mapping; instead, “BJ-derived cell line” was selected as the final output. However, even when the correct ontology term was present among the top 50 candidates, GPT-4o did not always select it. For dataset GSE44157, the “murine mesenchymal stem cells” cell line was correctly identified in the input text and the appropriate ontology term (“mesenchymal stem cell line”) was ranked 13th based on the embedding space similarity search, yet GPT-4o selected the specific mesenchymal cell line “RCB1991” despite the lack of supporting evidence.

## Discussion

In this study, we systematically evaluated the performance of GPT-4o for entity-to-ontology annotation, focusing on two important sample descriptors commonly seen in transcriptomic experiments: mouse strains and cell lines. Using 9,390 manually curated experiments from Gemma and 5,230 associated publications across the two annotation tasks, we show that GPT-4o can accurately identify and normalize the majority of relevant entities.

GPT-4o performed particularly well on the mouse strain annotation task, achieving correct annotation for 77% of experiments and high average recall and precision. Considering that the input text - consisting of the GEO experiment metadata and, when available, sections of the associated paper - was long, heterogeneous, and inconsistent, often containing discrepancies or typographic errors, we view this result as very promising. In contrast, the straightforward string-matching approach, while achieving relatively high recall, exhibited extremely poor precision and correctly annotated only 6% of experiments. This highlights the limitations of methods based solely on lexical similarity and the advantages of LLMs capable of contextual reasoning when interpreting free-text experimental descriptions.

We observed lower performance on the cell line annotation task, where 59% of experiments were annotated correctly. We believe this is partially due to the much larger Cell Line Ontology (CLO) dictionary, consisting of over 46,000 terms (compared to 156 mouse strain ontology terms), which was too large to be directly supplied to the LLM as input. Instead, we had to apply a workaround, utilizing OpenAI’s text-embedding model and similarity search over embedding vectors to provide GPT-4o with candidate cell line terms for annotation. Although this approach allowed us to implement the retrieval-augmented generation (RAG) framework, it also introduced errors arising from the retrieval and ranking of candidate ontology terms. It is possible that models that can fit the cell line terms (∼2.8M tokens) in the context window would alleviate this issue.

Additionally, CLO terms are known to have high lexical complexity, frequently consisting of short, similar alphanumeric codes that pose a challenge even for human curators. In line with this, Riquelme-Garcia et al. (2025), who evaluated various types of GPT models on a biological sample label annotation task, reported the lowest performance for CLO compared with the three other ontologies.

We found that in many cases GPT-4o’s errors closely resembled those made by human curators. They often arose from typographical mistakes in the GEO record, inconsistent naming or discrepancies between GEO metadata and the associated publications. Consistent with this observation, GPT-4o was more likely to make mistakes in experiments where string matching also failed to detect the correct mouse strains. We also observed occasional hallucinated annotations, where GPT-4o produced ontology terms not explicitly mentioned in the input text.

Importantly, however, the supporting quotes from the input text returned by the model were reliably accurate, allowing errors to be readily identified during review.

Interestingly, GPT-4o identified over 200 errors in Gemma’s annotations, even though the original curation was performed manually by trained data curators using assistance tools for mapping free text entries to ontology terms and following strict guidelines. In addition, these annotations are often reviewed by a second curator and occasionally subjected to systematic quality audits. These errors often stemmed from inconsistent reporting within GEO metadata and/or the associated publications, problems that are very time-consuming and difficult for humans to catch. GPT-4o was apparently able to overcome this by analyzing the entirety of the provided information.

Our results on LLM performance in the entity-to-ontology annotation task are more favorable than those reported in recent literature. Riquelme-García et al. (2025), who applied GPT-based models to automatically assign terms from multiple ontologies to biological sample labels, found that without fine-tuning, the base models performed poorly, with precision across models not reaching 20%. Because the LLMs were given only sample labels as input, the lack of broader context may have limited their ability to select the correct ontology identifier. In addition, since a RAG framework was not used in the study, the models often produced random identifiers that did not correspond to existing ontology terms. Kainer et al. (2025) investigated the ability of GPT-4o to annotate plant phenotype observations with ontology terms using several workflows, both with and without the RAG framework. Their RAG-based workflows - “descriptor to embedding” and “descriptor to concept to embedding”, which most closely resemble our mouse strain and cell line annotation approaches, respectively - achieved a highest recall of 0.517 and a highest precision of 0.397. A possible explanation for the lower performance in their study is the greater complexity of the annotation task, which required identifying and mapping multi-word phenotype concepts rather than single-entity mentions; however, at the same time, the model was given only single-sentence inputs, limiting the volume of text to be analyzed.

Our study has several limitations. First, we evaluated only a single LLM model. At the time of the study, GPT-4o was the most advanced GPT model and a successor to GPT-4, which was reported to outperform earlier GPT models, as well as some open-source LLMs, in several relevant studies (Aldeen et al., 2023; Chen et al., 2025; Groza et al., 2024; Poretsky et al., 2025). It is possible that newer models will give improved performance or pose additional challenges. Second, although Gemma’s transcriptomic experiments are carefully curated, the annotations do not constitute a definitive ground truth, and in fact, even GEO records and publications should be treated with caution. This is illustrated by the fact that Gemma contains curation errors, many of which were propagated from the sources. Although these were corrected before calculating accuracy measures for GPT-4o, some disagreements between GPT-4o and the reference annotations may still reflect limitations of the reference data rather than model errors. Third, for the cell line annotation task, performance depended on the embedding-based retrieval of candidate ontology terms. Failures to retrieve the correct candidate prevented correct ontology normalization even when the entity mention was correctly identified in the first stage, meaning the measured accuracy does not reflect the performance of the language model alone. Finally, we did not measure inter-annotator agreement among human curators, and therefore cannot directly compare model performance to the natural variability present in manual curation.

Overall, our findings indicate that GPT-4o, and likely other currently available LLM models, cannot fully replace a human curator for the entity-to-ontology annotation task. However, they show clear promise as assistive tools that can improve both the efficiency and quality of annotation. An effective curation workflow would adopt a human-in-the-loop model, in which LLMs perform initial screening of input text, identify and normalize entities or concepts, and provide results together with the supporting quotes for validation by human curators. Such an approach could help scale curation efforts for rapidly growing biomedical databases and repositories while preserving human oversight. Future work could explore tighter integration of LLMs into interactive curation workflows, the potential benefit of fine-tuning, improved retrieval strategies for large ontologies, and extension of this approach to additional metadata categories and experimental modalities.

## Author contributions

PP: Designed the study, directed work, contributed to manuscript development. SR: Contributed to study design, directed work, drafted the manuscript. BOM: Contributed to study design, performed experiments and evaluation. BX, AX, CY: Manual reviewed discrepant annotations, conducted Gemma re-curation.

## Acknowledgements

We thank current and former Gemma curators who manually annotated thousands of GEO experiments. We also thank GEO and thousands of GEO data submitters for providing a rich source of data. Finally, we acknowledge the efforts of the developers and maintainers of the ontologies used in this study.

## Code and data availability

The spreadsheets listing Gemma’s experiments used for evaluation of mouse strain and cell line annotation tasks, along with manual and predicted annotations, GPT-4o-generated supporting quotes, ontology URIs, performance metrics, curators’ notes and other details are available at https://github.com/PavlidisLab/GPT_annotate. The repository also contains GPT-4o prompts and R scripts for running the experiments and evaluating the models’ performance.

